# Climate change decreases the likelihood of tropical forest community persistence with a strong mediation of plant-plant network structures

**DOI:** 10.1101/2023.12.20.572696

**Authors:** Zijing Luo, Hanlun Liu, Yuanzhi Li, Weimin Wang, Margaret M. Mayfield, Chengjin Chu

**Affiliations:** State Key Laboratory of Biocontrol, School of Life Sciences, Sun Yat-sen University, 510275 Guangzhou, China; School of BioSciences, University of Melbourne, Parkville, 3010 Victoria, Australia; State Key Laboratory of Biocontrol, School of Ecology, Shenzhen Campus of Sun Yat-sen University, 518107 Shenzhen, China; Shenzhen Ecological and Environmental Monitoring Center of Guangdong Province, 518049 Shenzhen, China; Guangdong Greater Bay Area, Change and Comprehensive Treatment of Regional Ecology and Environment, National Observation and Research Station, 518049 Shenzhen, China; State Environmental Protection Scientific Observation and Research Station for Ecology and Environment of Rapid Urbanization Region, 518049 Shenzhen, China

**Keywords:** Barro Colorado Island, community persistence, ecological network, network structure, structural stability, temporal dynamic

## Abstract

Climate change is known to negatively impact tropical forests; yet how climate change impacts tree community persistence at local scales remains less clear. Using data from a long-term tropical forest census plot over 25 years, we constructed plant-plant interaction networks based on tree growth. We then quantified community persistence as feasibility domain of constituent species using recently developed frameworks of structural stability. We found a decrease in structural stability under warming and precipitation changes over time as evidenced by both direct environmental effects and indirect effects via network structure; and indirect effects were stronger than direct effects. Among these structures, facilitation:competition degree ratio and competitive transitivity were most positively related with structural stability. Our study highlights how the lens of plant-plant interaction networks can identify novel details about risk to tropical forest diversity under climate change at local scales. Insights from this work will be helpful for aligning forest management activities with areas under the greatest risk of species loss.

## INTRODUCTION

Climate change is one of the major factors threatening forest communities (Dawson et al., 2011; Hickling et al., 2006; Parmesan, 2006). Climate changes such as warming and drought cause disruptions in tree demography (Wright, 2005; Teller et al., 2016). Warming increases tree growth and decreases survival and recruitment, while drought reduces the response of tree growth to warming (Briffa et al., 1998; Doak & Morris, 2010; Morris et al., 2020; Swetnam & Betancourt, 1998). In addition to altering demographic rates of individual species, climate change may also impact interactions among species (Alexander et al., 2015; Alon & Sternberg, 2019; Esch et al., 2018) and the outcome of species coexistence (Van Dyke et al., 2022; Wainwright et al., 2018). For example, experimental precipitation change doesn’t only affect plant vital rates and pairwise interactions, but also affects stabilizing niche and fitness differences, which in turn determines if coexistence occurs (Van Dyke et al., 2022; Wainwright et al., 2018). Unlike experimental pairwise systems, diverse natural forest systems are complex and thus it is unclear if the impacts of climate change observed in experimental studies of pairs of plant species (most are annual plants) reflect generalizable consequence of climate change that hold across entire networks of interacting species.

Previous estimations of interaction networks usually have some critical weaknesses. These include: (1) using correlations of co-occurrences as proxies for interactions without considering individual performance (Blanchet et al., 2020; Losapio et al., 2021), (2) avoiding hard-to-measure interactions such as plant-plant interactions (Bimler et al., 2023; Simmons et al., 2018), and (3) majorly considering one type of interaction (either mutualism or antagonism) and neglecting the continuum of mixed interaction types (Mougi & Kondoh, 2012; Montesinos-Navarro et al., 2017) or the mixed roles certain species have across a whole community (Genrich et al., 2017). Other difficulties are evident in studies attempting to quantify multispecies coexistence (Barabás & Andrea, 2016). For instance, classic indices of network stability usually only allow for a single interaction type (e.g. topological robustness; Barabás et al., 2014; Meszéna et al., 2006), or rely on estimations of all parameters in a population model (e.g. local stability; Aufderheide et al., 2013) including intrinsic demographic rates which are difficult to be quantified in long-lived plants, like trees. Recent advances in structuralist networking approaches (Rohr et al., 2014; Saavedra et al., 2017; Song et al., 2018), however, provide an alternative to evaluating possibilities of multispecies coexistence only based on species interaction matrix. Structural stability from these approaches quantifies the parameter space of intrinsic demographic rates that allows species to persist (i.e. species all have positive abundances at equilibrium) with a given interaction matrix. Since intrinsic demographic rates are thought to depend on environmental contexts, structural stability can represent community persistence over time and across environmental gradients (Rohr et al., 2014; Saavedra et al, 2017; Song et al., 2018). What’s more, calculation of structural stability can be applied on matrices with both positive and negative interactions increasing the biological realism used to study horizontal network stability. These two advantages make the approach to structural stability modelling suitable for measuring the likelihood of coexistence in complex forest communities.

In calculating structural stability, it is important to consider how species interactions are organized, i.e., interaction network structures and potential factors influencing structural stability (Bascompte & Jordano, 2014; Trøjelsgaard & Olesen, 2016). Interaction network structures may be important in mediating effects of climate change on structural stability in plant-plant networks, as have been shown in both plant-pollinator and predator-prey networks. Studies of cross-trophic networks have found that structural stability was influenced by turnover in seasonal species interactions and changes in patterns of network nestedness (Rohr et al., 2014; Saavedra et al., 2016a; Saavedra et al., 2016b). In a theoretical study, Saavedra et al. (2017) found that though there is no pairwise coexistence predicted, the structural stability analysis reveals that coexistence of three plant species is still feasible, suggesting competitive intransitivity can result in multispecies coexistence. Other studies have found that network structures may link other aspects of network stability to specific environment conditions. For example, in plant-pollinator systems, climate variation is known to affect network modularity and nestedness, which in turn influences community ‘robustness’ (Bascompte et al., 2019; Dalgaard et al., 2013; Liu et al., 2021). The same pattern has been demonstrated in host-parasitoid systems, where habitat fragmentation affects modularity and then makes communities susceptible to species loss (Grass et al., 2018). Given these previous studies it seems possible that network structure may also reveal how climate changes alter the structural stability of complex plant communities as described by plant-plant interaction networks. Big gaps in our theoretical expectations of these relationship, however, include what these effects are and how they might vary across spatial scales. What we know from classic ecological theory suggests that species interactions and/or local environmental factors (e.g. topography) may contribute more to diversity maintenance at small scales, while other processes like dispersal may be more prominent at large scales (Ricklefs 2003; Ricklefs & He, 2016). Studies of these predictions with real data, however, are lacking. One major reason for this is the lack of data on species interactions and coexistence across spatial scales over time. Only time series data can allow us to test predictions on how well temporal fluctuations in structural stability at short time scales reflect long-time dynamics.

In this study, we aim to fill these knowledge gaps using plant performance and time-series data from the long-term forest plots on Barro Colorado Island (BCI), Panama. Specifically, we ask the following three questions: 1) How does the likelihood of species coexistence in tropical forests change under climate change over time? 2) What dose network structure tell us about tree coexistence under different topographies and climate conditions? And 3) how does the likelihood of coexistence change with network structure, climate factors and local environmental conditions across different spatial scales? To answer these questions, we used BCI forest data to estimate tree species interaction networks based on individual tree growth, which we then used to calculate network structures and stability. We hypothesized that the likelihood of tree coexistence in this forest system varies under climate change, and is also determined by different local topography conditions. These environmental factors might impact the likelihood of tree coexistence directly as well as indirectly via network structure. We also hypothesized that different spatial scales have different determining factors of tree coexistence among interaction structures and environmental conditions.

## MATERIALS AND METHODS

### Study site

The 50-ha permanent forest plot is located in Barro Colorado Island (BCI), Panama (9°9′ N, 79°51′ W), with the elevation ranges from 120-160 m, annual mean temperature of 27℃ and annual rainfall of 2600 mm (Dietrich et al., 1992). BCI plot is the representative plot of Forest Global Earth Observatories network (ForestGEO, http://www.forestgeo.si.edu/; Davies et al., 2021) with the longest investigation time. Established in 1980 (Condit, 1998), BCI plot was first censused in 1981-1983, re-censused in 1985 and kept censusing every 5 years since then. All freestanding woody stems with diameter at breast height (DBH) of at least 1 cm were mapped, tagged, identified and measured. Aimed at temporal variation of interaction networks, we used data of six censuses from 1990 to 2015 (Data available from the Dryad Digital Repository: https://doi.org/10.15146/5xcp-0d46; Condit et al., 2019), where species with abundance more than 500 were included. We did not use the early censused data in order to make the consistence of methods in measuring tree growth (Condit, 1998). The plot was divided into 10 m × 10 m, 20 m × 20 m and 50 m × 50 m quadrats, aimed to verify whether the relationships among variables would be consistent at different spatial scales or not.

### Environmental variables

We considered spatial and temporal variations of environmental factors, which were measured by topography and climate variables separately. Elevation was defined as mean elevation of four corners of a significant quadrat. Elevation data were recorded for each quadrat of 20 m × 20 m scale as primary topography information, based on it we calculated other topological variables including slope, convexity and aspect (Baldeck et al., 2013; Harms et al., 2001), which could be estimated for larger (50 m × 50 m) and finer (10 m × 10 m) scales. Slope was calculated as mean slope of planes defined by any three corners of a quadrat. Convexity was calculated by elevation of a quadrat minus average elevation of its surrounding quadrats. Aspect means the direction where the slope faced, and then be calculated as sin(aspect) and cos(aspect) representing different directions (Legendre et al., 2009).

Climate variables here were derived for each census year, with each year containing four variables that were mainly found to be important in influencing species interactions and biodiversity in previous studies (Bellard et al., 2012; Blois et al., 2013; Schleuning et al., 2016): mean annual temperature (MAT) calculated by the mean of total maximal and minimum temperature of 12 months, annual precipitation (AP) calculated by the sum precipitation of 12 months, temperature seasonality (TS) calculated by 100 times the standard deviation between seasons, and precipitation seasonality (PS) calculated by coefficient of variation between seasons. Primary monthly climate data were downloaded from WorldClim 2.1 (https://worldclim.org/data/monthlywth.html; Harris et al., 2020; Fick and Hijmans, 2017) at a spatial resolution of 2.5 minutes (∼21 km^2^).

### Plant-plant interaction network estimation

Individual growth was calculated by the difference in DBH of tree individuals between two neighboring censuses, and then was fitted in individual growth model to calculate interaction coefficients. We used the model from Li et al. (2021), assuming that interaction effects were positively related with DBH of neighbors while negatively related with distances of interacting species. The model was:

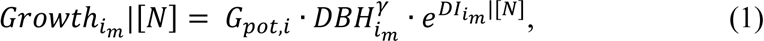

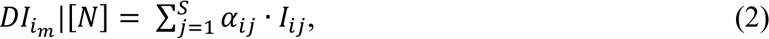

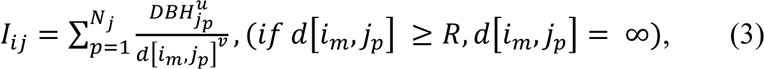

where *Growth*_*im*_|[*N*] represents growth of individual *m* of species *i* with *N* neighbours, calculated by the difference of DBH of a same individual tree between two consecutive censuses. *G*_*pot,i*_ represents the potential growth of species *i* without neighboring interactions. *DBH*_*im*_ and *DBH*_*jp*_ represent DBH of individual *m* of species *i* and individual *p* of species *j* separately. *S* means richness of neighbor species of a certain species *i*. α_*ij*_ means interaction coefficients. *d*[*i*_*m*_, *j*_*p*_] means distance of individual *m* of species *i* and its neighbor individual *p* of species *j*. *U* and *v* determined the effect of DBH and distance, here we followed the parameters set in Li et al. (2021) and set *u* = *v* = 1 in our study. *R* means the maximal distance where interaction could happen, here we set 10 m, 20 m and 50 m. Let *I*_*ij*_ represent the sum of DBH of species *j* near species *i* weighted by distance between individual *m* of species *i* and all its neighboring individuals of species *j* within radius *R* interaction effects. After logarithmic transformation of formula (1), we use *DBH*_*im*_ and *I*_*ij*_ as predictors to fit the individual growth with linear regression model, and the slopes of *I*_*ij*_ were interaction coefficients α_*ij*_. Since interactions between plants actually happen within neighborhoods, we divided the whole sample plot into quadrats of 10 m × 10 m, 20 m × 20 m and 50 m × 50 m scales following the values of *R* above. For each quadrat, we established local plant-plant interaction network expressed as *n* × *n* matrix (*n* means *n* species appeared in this quadrat) with interaction coefficients α_*ij*_as elements.

### Network structures and structural stability

We used the number of species in each quadrat as their network size. As there were different types of interactions among plants, we extracted facilitation network and competition network from each local interaction network by the sign of α_*ij*_, then calculated the ratio of the numbers of positive and negative coefficients (facilitation:competition) as the proportion of the two types of networks. For each type of network, we calculated its modularity, that refers to the level of compartmentalization of a network where the number and strength of interactions inside modules are larger than that of the outside, and nestedness, which means the extent that species interacting with specialists is the subset of those with generalists.

And we then calculated intransitivity (that happens when competitors are limited by one another) for competition networks, with higher competitive intransitivity indicating more nonhierarchical competitions (e.g. rock-paper-scissors) in networks.

To calculated modularity, we used the R package ‘Infomap’ (Maps of information flow; Rosvall & Carl, 2008), considering the directions and strengths of interactions among plant species specifically. This method calculated modularity through the minimum amount of information *L* the random walker needed (Farage et al., 2021), and *L* was negatively related to modularity. We used the parameter of WNODF (Weighted Nestedness metric based on Overlap and Decreasing Fill; Almeida-Neto & Ulrich, 2011) in the R package ‘bipartite’ to calculate nestedness, through quantifying the overlap and decreasing of interactions in networks, and WNODF was positively related with nestedness. Since both modularity and nestedness were sensitive to network size and connectivity, we used null model ‘Quantitative Swap and Shuffle Model’ in the ‘vegan’ package to standardize them. This null model reshuffled positions of elements in interaction network matrix 1000 times, and the value obtained was used to calculate z-score with formula 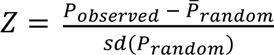. Intransitivity was calculated using Laird & Schamp’s method (2006). Firstly, we transformed interaction matrix into competition outcome matrix through setting 1 or 0 for winner or loser in pairwise competition by comparing absolute value of α_*ij*_ and α_*ji*_. Then the variance of new matrix represented intransitivity. The higher the value of variance, the lower the intransitivity.

We used individual growth as a proxy of fitness, specifically reflecting the DBH size class structure of community. The stability of growth-based models essentially represents the relative balance of community size class structure, which prevents the occurrence of species with much faster growth than other species. Since accumulating advantage of fast-growing species to slow-growing species may translate into potential competitive exclusion during subsequent life stages such as reproduction, the growth-based stability can serve as a proxy of tree species coexistence.

Structural stability was used to measure likelihood of coexistence of species in local interaction networks. Describing the parameter space of intrinsic growth vector of all species in network which was called feasibility domain, structural stability quantified this feasibility domain by calculating the possibility that all species had positive abundance at equilibrium. Therefore, structural stability represented the overall tolerance range of a community to specific fluctuating environment, and the larger the parameter space, the higher the likelihood of species coexistence (Rohr et al., 2014; Saavedra et al., 2017; Song et al., 2018). Its mathematical definition is given in Song et al. (2018).

### Statistical analyses

Linear regression model was used to fit the relationships between structural stability and environmental factors, including climate factors (MAT, AP, TS and PS) and topography variables (elevation, slope, convexity and aspect). We then used Principal Component Analysis (PCA) to reduce dimensions of variables. For topography variables and network structure variables, we extracted the first two axes separately, where PC1 and PC2 were then used as new variables in the following analyses. The first two axes of network structure PCA together explained nearly half variation of network structure, with 48% at the small spatial scale (10 m × 10 m) and 54% at the medium spatial scale (20 m × 20 m), while explained most structural variation (88%) at the larger spatial scale (50 m × 50 m). At the 10 m × 10 m scale, the first PC (Str_PC1_) was mainly related to network size, nestedness of facilitation network and modularity, nestedness and intransitivity of competition network. The second PC (Str_PC2_) represented the degree ratio between facilitation and competition network, nestedness of facilitation network and intransitivity of competition network. Results were almost the same for the 20 m × 20 m scale. What was different at the 50 m × 50 m scale was that Str_PC1_ explained variation of all seven structural variables, while Str_PC2_ only represented large network size and small nestedness of competition network. Specific values of loadings were shown in Table 1. For topography PCA, the first two axes together explained nearly half variation of topography, which were continuous among all three spatial scales (49 % in 10 m × 10 m; 52 % in 20 m × 20 m; 56 % in 50 m × 50 m). The first axis (Topo_PC1_) represented environment with high elevation, low slope and convexity, and the second axis (Topo_PC2_) represented environment with low sin(aspect) and cos(aspect).

**TABLE 1.** Loadings of the first two axes of network structure PCA at three spatial scales.

|  | 10 m × 10 m |  | 20 m × 20 m |  | 50 m × 50 m |  |
| --- | --- | --- | --- | --- | --- | --- |
|  | PC1 | PC2 | PC1 | PC2 | PC1 | PC2 |
| NS | 0.85 | 0.08 | 0.83 | -0.32 | 0.47 | 0.82 |
| DR | 0.16 | 0.82 | 0.73 | 0.43 | 0.89 | 0.09 |
| FNM | 0.04 | -0.07 | -0.18 | 0.23 | -0.98 | 0.05 |
| CNM | 0.48 | 0.13 | 0.47 | 0.48 | -0.97 | 0.10 |
| FNN | 0.34 | 0.52 | 0.56 | 0.06 | 0.92 | 0.12 |
| CNN | -0.83 | 0.18 | -0.67 | 0.45 | 0.65 | -0.62 |
| CNI | -0.46 | 0.60 | 0.04 | 0.88 | -0.92 | 0.03 |

NS: network size; DR: degree ratio of facilitation and competition network; FNM: facilitation network modularity; CNM: competition network modularity; FNN: facilitation network nestedness; CNN: competition network nestedness; CNI: competition network transitivity. Note that the higher the FNM, CNM and CNI, the lower the corresponding modularity and intransitivity.

We finally used Structural Equation Model (SEM) to explore direct and indirect effects among variables, with the ‘piecewiseSEM’ package in R (Lefcheck et al., 2020; Lefcheck, 2016). The primary model included simple linear models of structural stability explained by all other variables, and network structure PCs explained by topography PCs and climate variables. We removed non-significant paths with P > 0.05 step-by-step, and the goodness-of-fit of final model was measured by Fisher’s *C*-test with P > 0.05. We conducted SEMs for each climate factor separately because of the limitation on climatic samples (here we included five samples corresponding to each census year for each climate factor). During the process of selecting SEMs, Topo_PC2_ did not significantly influence other variables, therefore we moved it away and changed the rest topography PC (Topo_PC1_) into its representative primary variables (i.e. elevation, slope and convexity). All statistical analyses were conducted using R 4.1.1 (R Core Team, 2021).

## RESULTS

### Temporal variation of community persistence under climate change

Structural stability of feasibility domain decreased temporally from 1990 to 2015, while temperature increased over the same time period (Figure 1a). This reflects a pattern that structural stability decreased towards high annual mean temperature (Figure 1a, S2). These results were consistent across all three spatial scales (10 m × 10 m, 20 m × 20 m and 50 m × 50 m). The larger the spatial scale, the steeper the decreasing trend of structural stability over time (Figure 1a).

**FIGURE 1.**
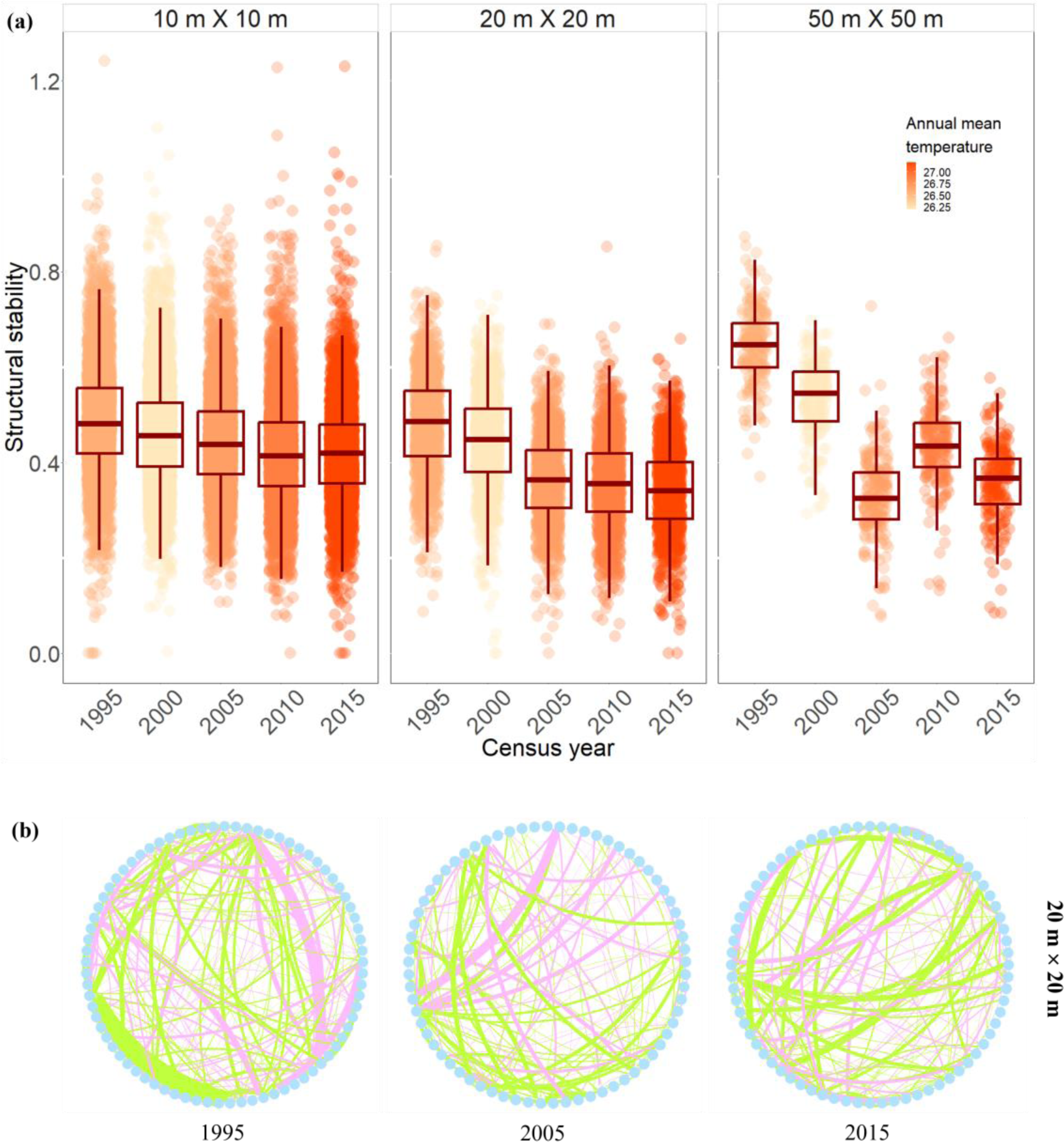
Dynamics of structural stability of the BCI plot in 1990-2015 census years (a) and species interaction networks (b). (a) Boxplot of each census year shows the maximum, top quartile, median, bottom quartile, and minimum value of structural stability. Structural stability of local interaction network was calculated at three spatial scales: 10 m × 10 m, 20 m × 20 m and 50 m × 50 m. (b) Tree species interaction networks (the spatial scale of 20 m × 20 m as an example) of three census years. Each blue dot represents a specific plant species, and the green and pink lines represent facilitation and competition interactions among tree species. The width of lines is associated with strengths of interactions.

Regarding to climatic factors, higher annual mean temperature, annual precipitation, and temperature seasonality were associated with lower structural stability at all three spatial scales (P < 0.05). Only precipitation seasonality showed the opposite pattern where higher precipitation seasonality was associated with higher structural stability at two of three spatial scales (P < 0.05) (Figure S1). These results provide evidence supporting a possible causative association between climate factors and declining structural stability.

### Mediation of network structures on tree coexistence in different environments

We found that both climate and topography variables had indirect effects on structural stability at most spatial scales. For climate variables, the indirect effects via network structures were stronger than direct effects in five of six SEMs of different climate variables and spatial scales (Figure 2). This is evidenced by strong impacts of network structures on structural stability. Specifically, annual mean temperature had stronger indirect effects (-0.10) than direct effects (-0.06) at the 10 m × 10 m scale (Figure 2a), and at the 20 m x 20 m scale (indirect effect (-0.20) and direct effect (-0.15, respectively; Figure 2e). For annual precipitation, indirect effects were stronger than direct effects at all three spatial scales (Figure 2b, f, j). These results were similar for temperature seasonality and precipitation seasonality (Figure 2c-d, g-h, k-l). On the other hand, the direct paths between topography variables and structural stability were weaker than for climate factors (Figure 2; Figure S2). The indirect effects of different topography variables were stronger than their direct effects in half of all SEMs across spatial scales (two of six SEMs for elevation, three of six SEMs for slope and four of six SEMs for convexity).

**FIGURE 2.**
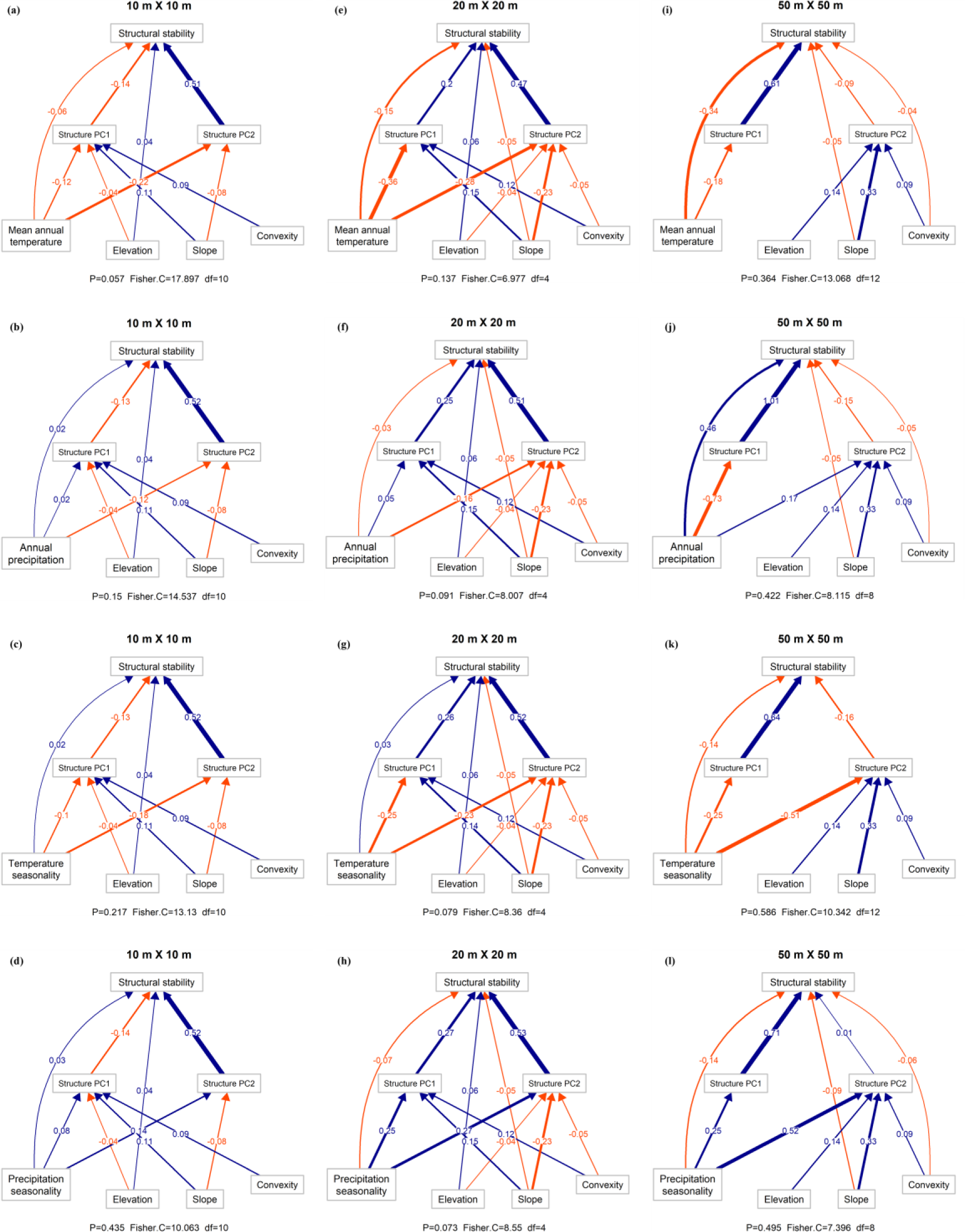
Structural Equation Models of relationships among climate factors, network structure PCs (represented by seven network structure variables) and structural stability. Climate factors were represented by mean annual temperature (a, e, i), annual precipitation (b, f, j), temperature seasonality (c, g, k) and precipitation seasonality (d, h, l) separately. Three spatial scales were calculated including 10 m × 10 m (a-d), 20 m × 20 m (e-h) and 50 m × 50 m (i-l). Arrows represent significant paths (excluding two paths from Str_PC_ to structural stability at the 50 m × 50 m scale; here we retained them with the consideration of ecological significances), with standardized path coefficients represented by labels on them. Different colors of paths represent different types of effects between variables, where blue is for positive interactions and red for negative interactions.

For different kinds of environmental variables, our results found that climate factors had stronger effects on structural stability than topography factors both directly and indirectly at most spatial scales. These generally stronger effects of climate factors were showed in the stronger direct (29 of 36 comparisons) and indirect (34 of 36 comparisons) effects of four climate factors to structural stability than three topography factors at three spatial scales (Figure 2). Additionally, among these climate factors, our results further revealed that annual trends of temperature and precipitation had stronger effects on structural stability than temperature and precipitation seasonality. This was illustrated by the comparisons between annual trend with seasonality of temperature and precipitation respectively, where annual trends had stronger direct (five of six comparisons) and indirect (four of six comparisons) effects on structural stability than seasonality generally (Figure 2).

Among these interaction structures, several played key roles in affecting community persistence. Specifically, facilitation:competition degree ratio and competitive transitivity showed the similar decreasing trends as structural stability did over the same time period (Figure 3a-b). These two interaction structures were significantly positively associated with structural stability (P < 0.01) (Figure 3c-h).

**FIGURE 3.**
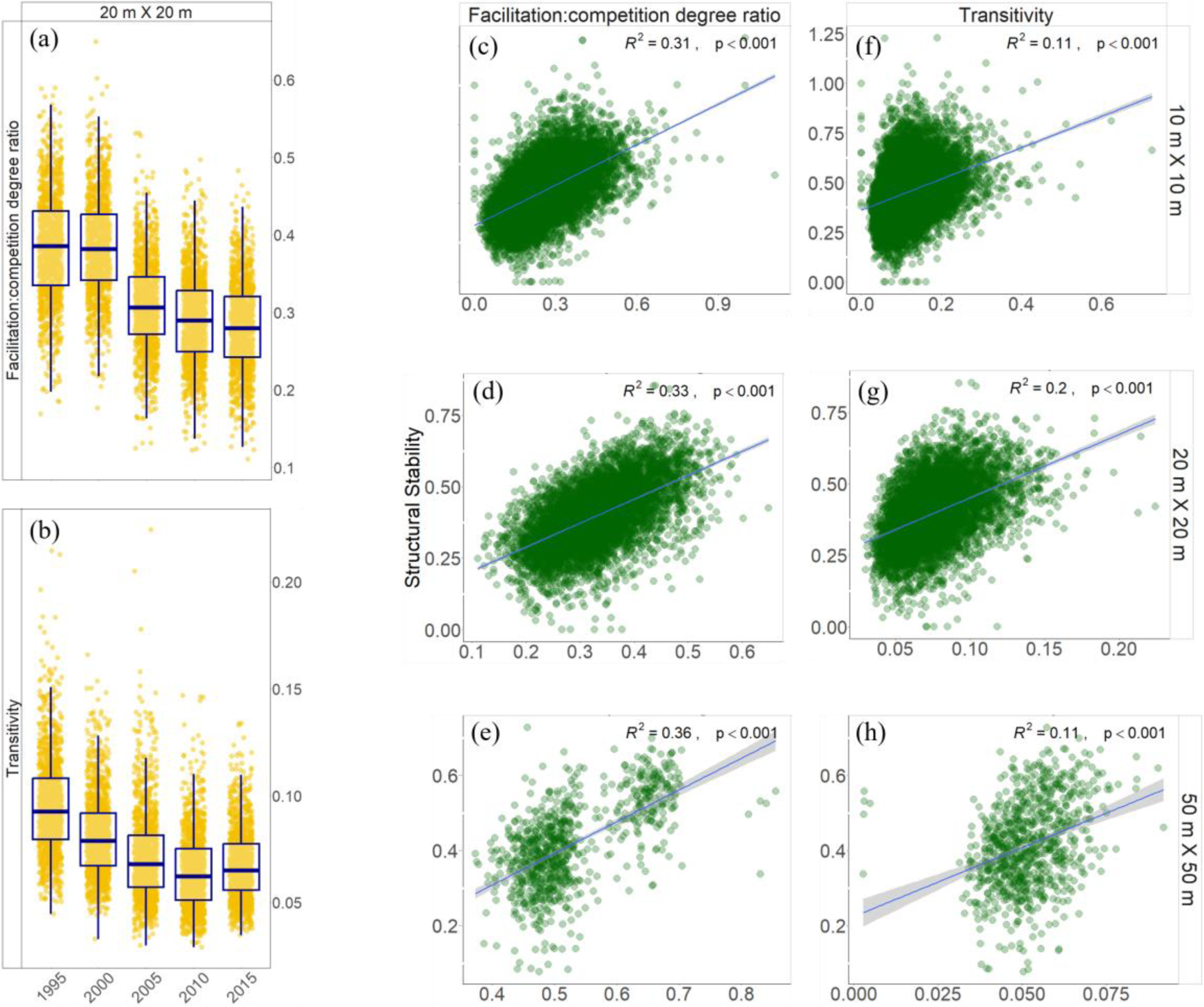
Dynamics of interaction structure of the BCI plot in 1990-2015 census years (a-b) and relationships between structures and structural stability (c-h). (a-b) Boxplot of each census year shows the maximum, top quartile, median, bottom quartile, and minimum value of interaction structures including facilitation:competition degree ratio and transitivity at 20 m × 20 m spatial scales. (c-h) Linear regressions of facilitation:competition degree ratio (c-e), transitivity (f-h) and structural stability were conducted for each panel. Relationships were calculated at three spatial scales: 10 m × 10 m (c, f), 20 m × 20 m (d, g) and 50 m × 50 m (e, h). The blue line in each panel is the regression trend line and grey shading represents the 95% confidence interval.

These results suggest that facilitative interactions and competition hierarchy might be critical to help forest community buffer effect of climate change on persistence.

### Effects of network structures and environments on community persistence across spatial scales

There were consistencies of the effects from network structures, climate factors and topography to structural stability across spatial scales. Firstly, there were significant and strong effects of network structure PCs on structural stability at all three spatial scales, with significant paths from Str_PC2_ to structural stability with standardized path coefficients of 0.51, 0.52, 0.52 and 0.52 at the 10 m × 10 m scale (Figure 2a-d), and 0.47, 0.51, 0.52 and 0.53 at the 20 m × 20 m scale (Figure 2e-h) respectively, which meant common network structures including ratio of facilitation versus competition, and competitive transitivity both promoted structural stability of networks at these spatial scales. These results were consistent when expanded to the large spatial scale, reflected by strong effects from Str_PC1_ to structural stability at the 50 m × 50 m scale (Figure 3i-l).

Secondly, the mediation effects of network structures were consistent and important across different spatial scales. At most spatial scales, indirect effects of climate factors on structural stability via network structures were stronger than direct effects. Specifically, at the small and medium spatial scales of 10 m × 10 m and 20 m × 20 m, effects of the annual trends and seasonality of temperature and precipitation on structural stability were all mainly indirect via network structures (Figure 2a-h). In contrast, when the spatial scale was expanded to 50 m × 50 m, there were two different situations in which annual trend and seasonality of temperature both affected structural stability directly (Figure 2i, k).

## DISCUSSION

Our study provides novel evidence that network structural stability in the BCI tropical forest plot decreased temporally during the past 25 years, a decrease strongly correlated with common climate factors reflective of global climate change, notably increasing temperatures. We found that indirect effects mediated by interaction network structures were more important than direct effects of environmental variables on structural stability at small and medium spatial scales. Specifically, facilitation:competition degree ratio and competitive transitivity were the most important interaction structures affecting structural stability. Thus, local interaction network structures were prominent in mediating the effects of climate change on multispecies coexistence, while at larger spatial scales, direct effects of climate factors may play more important roles in determining the likelihood of community persistence.

### Temporal variation of community persistence under climate change

The finding that temporal decreases in the likelihood of multispecies coexistence were common in the BCI plot (Figure 1a) is surprising and has potentially pressing implications for the conservation of these forest systems. Although many other aspects related to coexistence have been revealed to be impacted by climate change— including species extinction (Román-Palacios & Wiens, 2020) and richness (Descombers et al., 2020), niche overlap and fitness difference of species (Blackford 2020; Perez-Ramos 2019; Matias 2018), community productivity (Isbell et al., 2015), community biomass stability (Fussmann et al., 2014)—such hazard has not been demonstrated in plant community persistence, especially the obvious temporal trend of decreasing persistence revealed at the first time in our study. This decrease of the likelihood of community persistence might result from the impact on species climatic niches. Since species have their specific tolerance ranges of temperature and precipitation that are adaptive to their living condition (i.e. the climatic niche), fluctuations of climate conditions can cause species migration towards other suitable places. Then the geographical range of species would change (Soberón, 2007; Holt, 2009; Bellard et al., 2012). Therefore, species loss would happen at local scales, and geographical range of these migrating species might become smaller at larger spatial scales. These two aspects might reflect the potential hazard of climate changes on long-term persistence of these species.

Climate factors significantly influenced the likelihood of community persistence (Figure 2, S2). Annual mean temperature negatively influenced structural stability across all three spatial scales (Figure 2, S2), which might result from changes in individual growth rates induced by temperature fluctuation, and therefore further influences community structure (Lessard et al., 2011; Ratkowsky et al., 1982). Annual precipitation significantly affected species coexistence (Figure 2, S2), probably by changing individual growth (Lloret et al., 2016) and mortality (Niu et al., 2014; Thibault & Brown, 2008). Besides annual means, seasonality also affected structural stability significantly (Figure 2, S2). This might be caused by the fact that seasonality impacts the timing of species interaction and the variation of the interaction timing (Rudolf, 2019). Meanwhile, phenology also has potential impact on species coexistence by impacting temporal niche difference among species (Blackford et al., 2020).

### Direct and indirect effects between environments and community persistence

The mechanisms of temporal stability dynamics observed in our study involved both direct and indirect effects via network structures, though indirect effects were stronger (Figure 2). Results on indirect effects of environments fluctuations such as climate changes on stability were mediated by network structure, and this mediation is at the first time demonstrated through plant-plant interaction networks in the present study. This mediation of network structure is consistent with findings in other systems like plant-pollinator system, where local interaction network structure of modularity impacts wide range of coextinction behind primary extinction driven by environmental changes including climate, topography and insularity (Bascompte et al., 2019; Liu et al., 2021). This important mediation of interaction structures might result from the weak migration ability of plants. Under the limitation of movement, plants have to put more energy in gaining essential resources including lights and water in their local micro-environments (Bartels & Chen, 2010; Huston, 1979; Ricklefs, 1977), as a result, plants might become more susceptible to competition or facilitation from their neighboring plants. Furthermore, besides the interactions aboveground such as light competition, plant species in forest communities also link strongly with each other underground, which could be achieved through plant-fungi networks, where exchanges on resources like nutrients happened (Hart et al., 2003; Bennett et al., 2017). Thus, with above mechanisms, interaction structures certainly play key roles of mediation in plant-plant communities, leading to the significant indirect effects of fluctuating environments on multispecies coexistence.

Specifically, we found that facilitation degree proportion and competitive transitivity were the interaction structures most closely related with structural stability. Firstly, plants tend to positively interact with each other under environment threat. Since interactions between species are changeable under different environments, species usually corporate with each other to together resist harmful environmental condition while compete when resources are rich (Bertness & Callaway, 1994). Thus, as community persistence is decreasing under 25-year warming, plant species might tend to positively interact with each other, leading to the importance of facilitative interactions proportion to the whole community. Secondly, competitive transitivity in our study is positively related with structural stability, which seems contradicting with previous findings that intransitivity might promote coexistence. These previous studies mainly quantified interaction based on plant abundance (Ulrich et al., 2014; Soliveres et al., 2015). However, when estimation of interaction is more exact based on plant performance as we did, another study found that intransitivity is not as important as expected (Godoy et al., 2017).

On the other hand, environments also had direct effects on community persistence, which were reflected in both climate and topography factors (Figure 2), among which the direct effects might result from impacts of climate changes such as warming and drought that lead to species loss and further lead to following coextinction cascades. Direct effects of topography on multispecies coexistence might be realized through local micro-environments that determined the availability of light and soil moisture, level of soil pH (Pérez-Ramos et al., 2008; Perroni-Ventura et al., 2006), therefore directly impact growth of tree individuals and further indirectly impact resource competition among neighbouring plants. For further studies on temporal variation, data of other fitness components of plants or interaction partners of different trophic levels may be needed (Buzhdygan & Petermann, 2023), such as time series data (Deyle et al., 2016) of recruitment and reproduction of plants, or their herbivores, pollinators and seed dispersers by capturing their population dynamic in a much longer period of time.

### Relationships of biotic interactions, climatic factors and community persistence across spatial scales

The mediation effects of network structures were consistent when expanding to larger spatial scales in our study (Figure 2). However, another thing needs to be noticed is that, the mediation effects of network structures at larger spatial scales seemed not as important as that at smaller spatial scales. This may result from other ecological processes that have not been considered in our study, such as dispersal that influences species coexistence at larger spatial distance, which needs to be further examined.

Since network structures are calculated based on individual growth model, larger spatial distance among individuals may cause lack of explanation on interaction coefficients, leading to the probable less important mediation of network structures at large spatial scales.

Our results reveal the importance of spatial scale in multispecies coexistence, which can be taken into consideration in future studies, specifically including estimating species interactions based on species-specific interaction distances, for different species might have different spatial range for interaction in natural communities. What’s more, our results suggest the variations of the effects of interaction structures on coexistence across spatial scales, therefore, it is worth exploring the effects of interaction and dispersal on coexistence separately. Recently, this idea has already been explored in coastal ecosystem, where niche and dispersal have been disentangled to explore their effects on biodiversity maintenance (Loke & Chisholm, 2023).

In conclusion, quantifying persistence of forest communities remains a critical challenge for ecologists, and knowing how the likelihood of multispecies coexistence changes under changing environment has important implications for stability and biodiversity maintenance in forest communities. Here we focused on plant-plant interaction networks in a long-censused forest plot, and revealed that the likelihood of multispecies coexistence in this community decreased along census years, which was mainly driven by climate change especially climate warming, with the important and significant mediation effects of network structures. Coexistence based on individual growth model in our study assumed that the accumulated growth advantage of fast-growing species could be further translated to competitive advantage at subsequent life stages, which requires further experimental work to verify. In addition, specific ecological mechanisms underlying the plant-plant interactions (such as nutrients resource competition, facilitation by providing shades) and other (a)biotic factors determining community persistence are worth being explored, which could be based on developments of more refined data and methods, including performance data based on other fitness components, finer spatial-specific data, and more advanced methods to monitor dynamic of forest community. These further explorations could help ecologists broaden horizons to reveal mechanisms of community persistence in forest communities more deeply in a changing world.

## ACKNOWLEDGEMENTS

We thank Weitao Wang for the discussion on statistical analyses. This research was funded by the National Natural Science Foundation of China (31925027 and 32330064 to C.C.) and the National Key Research Development Program of China (#2022YFF0802300). H.L. and M.M.M were supported by the Australian Research Council (DP230100773).

## Supporting Information

**FIGURE S1.**
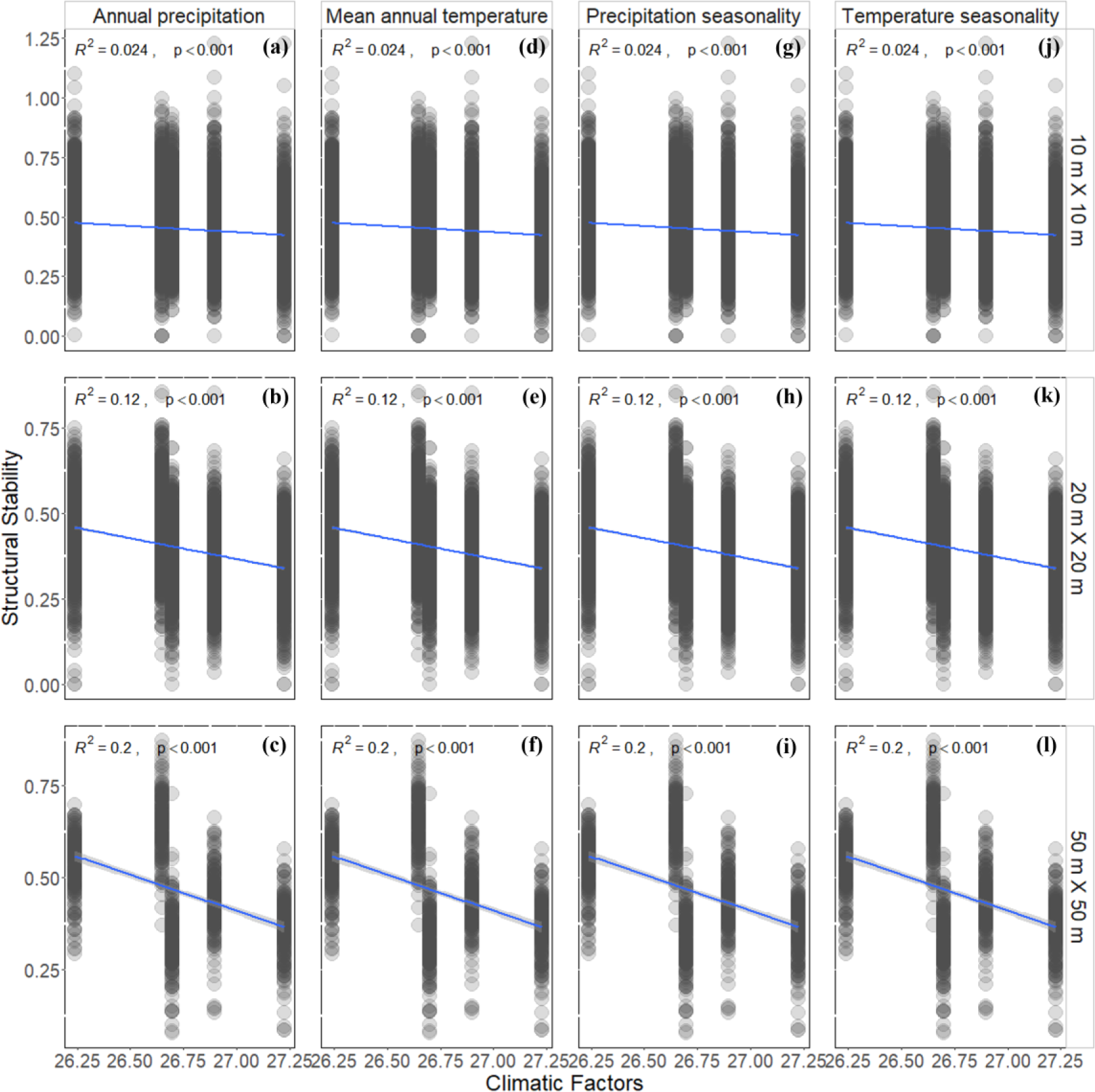
Relationships between climate variables and structural stability. Linear regressions of annual precipitation (a-c), mean annual temperature (d-f), precipitation seasonality (g-i), temperature seasonality (j-l) and structural stability were conducted for each panel. Relationships were calculated at three spatial scales: 10 m × 10 m (a, d, g, j), 20 m × 20 m (b, e, h, k) and 50 m × 50 m (c, f, i, l). The blue line in each panel is the regression trend line and grey shading represents the 95% confidence interval.

**FIGURE S2.**
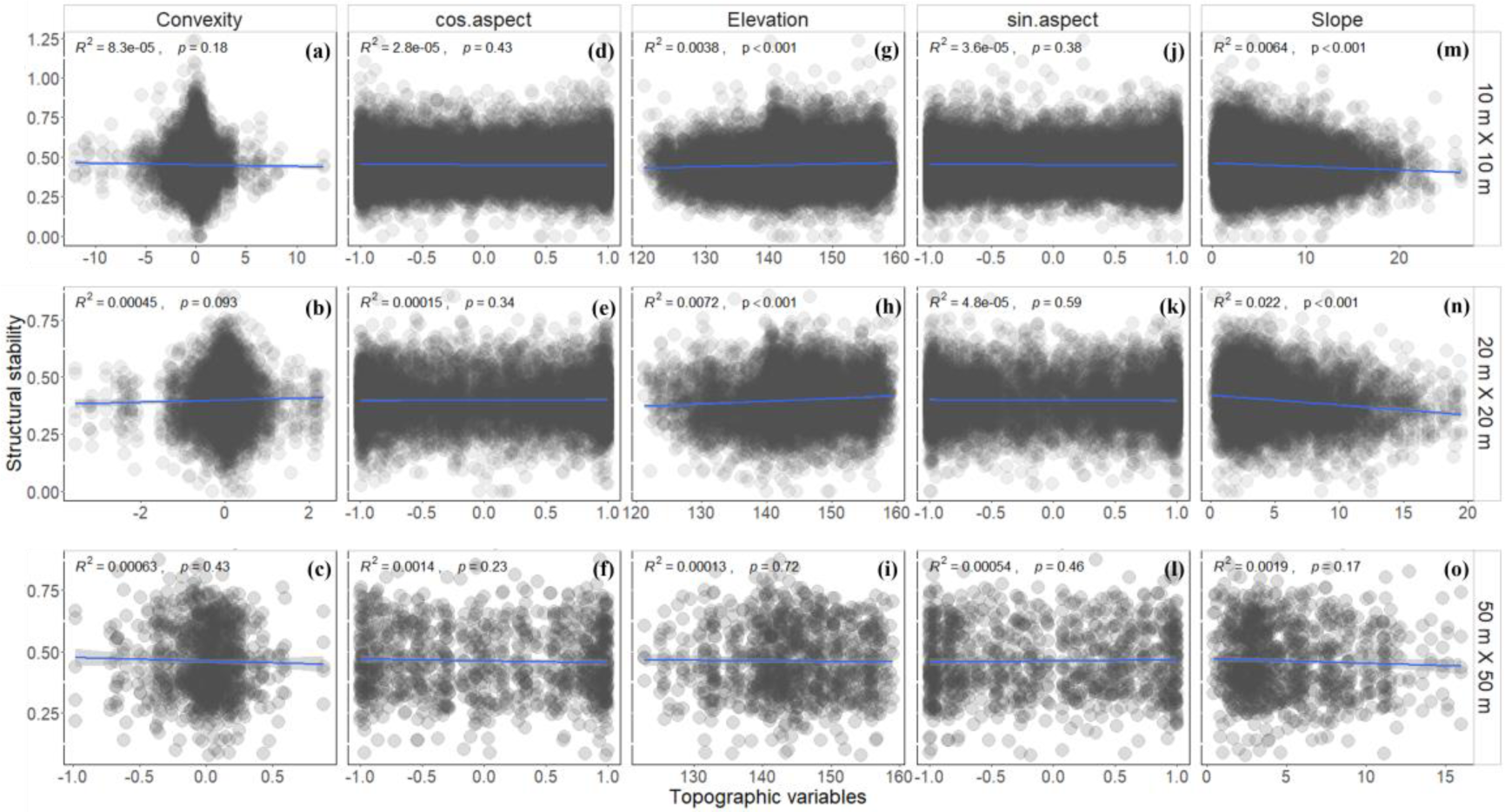
Relationships between topography variables and structural stability. Linear regressions of convexity (a-c), cos.aspect (d-f), elevation (g-i), sin.aspect (j-l), slope (m-o) and structural stability were conducted for each panel. Relationships were calculated at three spatial scales: 10 m × 10 m (a, d, g, j, m), 20 m × 20 m (b, e, h, k, n) and 50 m × 50 m (c, f, i, l, o). The blue line in each panel is the regression trend line and grey shading represents the 95% confidence interval.

